# Comprehensive Neural Representations of Naturalistic Stimuli through Multimodal Deep Learning

**DOI:** 10.1101/2025.04.15.646250

**Authors:** Mingxue Fu, Guoqiu Chen, Yijie Zhang, Mingzhe Zhang, Yin Wang

## Abstract

A central challenge in cognitive neuroscience is understanding how the brain represents and predicts complex, multimodal experiences in naturalistic settings. Traditional neural encoding models, often based on unimodal or static features, fall short in capturing the rich, dynamic structure of real-world cognition. Here, we address this challenge by introducing a video-text alignment encoding framework that predicts whole-brain neural responses by integrating visual and linguistic features across time. Using a state-of-the-art deep learning model (VALOR; Vision-Audio-Language Omni-peRception), we achieve more accurate and generalizable encoding than unimodal (AlexNet, WordNet) and static multimodal (CLIP) baselines. Beyond improving prediction, our model automatically maps cortical semantic spaces, aligning with human-annotated dimensions without requiring manual labeling. We further uncover a hierarchical predictive coding gradient, where different brain regions anticipate future events over distinct timescales—an organization that correlates with individual cognitive abilities. These findings provide new evidence that temporal multimodal integration is a core mechanism of real-world brain function. Our results demonstrate that deep learning models aligned with naturalistic stimuli can reveal ecologically valid neural mechanisms, offering a powerful, scalable approach for investigating perception, semantics, and prediction in the human brain. This framework advances naturalistic neuroimaging by bridging computational modeling and real-world cognition.

## Introduction

One of the central goals of neuroscience is to understand how the brain interprets and integrates information from the rich, dynamic, and multidimensional world we experience every day. Neural encoding models have proven valuable in this effort, offering quantitative predictions of brain activity under various conditions and providing insight into how information is represented in the brain^1–3^. Early work in this area focused on highly controlled, simplified stimuli to establish foundational principles. However, it is now widely recognized that such designs do not reflect the complex, continuous, and context-rich nature of real-world cognition. Rather than responding to isolated, static inputs, the brain continuously processes streams of multisensory information, integrates them over time, and interprets them within dynamic environments. This realization has motivated a growing push toward encoding models that better reflect the richness and variability of naturalistic experience ^4,5^.

Naturalistic paradigms—such as movie watching—have emerged as powerful tools for studying brain function in ecologically valid contexts^6,7^. Functional MRI (fMRI) during movie viewing increases participant engagement and enables researchers to investigate how the brain processes complex, time-varying inputs. Building on this foundation, several influential studies have modeled cortical responses to naturalistic stimuli using visual and semantic features ^5,8,9^, while others have advanced applications in non-invasive neural decoding^5,8,9^. These efforts have deepened our understanding of brain function in real-world settings. Yet, most existing encoding models are limited to a single modality, such as vision or language ^4,10–12^. Visual models (e.g., AlexNet^11^) perform well in early visual regions but generalize poorly to high-level areas, while semantic models (e.g., WordNet^13^) perform the opposite. Critically, semantic models often rely on dense human annotation. In early naturalistic encoding studies, trained raters watched the stimulus and labeled what was happening within each TR or short time window—for example, identifying objects, actions, or events present in the scene. These labels were then mapped onto a predefined semantic ontology (such as WordNet), yielding high-dimensional categorical feature vectors that served as regressors in encoding models. While this approach provides interpretable semantic features, it is labor-intensive, time-consuming, and inherently subjective, as annotations depend on rater judgment, labeling guidelines, and dataset-specific conventions, limiting scalability and reproducibility. These limitations highlight the need for automated, multimodal approaches that can more fully capture how the brain responds to naturalistic stimuli.

A second major challenge in naturalistic neuroimaging is improving model generalizability—the ability to predict brain responses to new, unseen stimuli. Many current models show sharp performance declines outside their training distribution. For example, a recent study reported that encoding models trained on one image set dropped to 20% accuracy when tested on out-of-distribution stimuli ^14^. Unimodal models may generalize within specific cortical domains but fail in broader, whole-brain contexts. To build truly useful encoding models, we must develop approaches that generalize across diverse stimuli, subjects, and cognitive domains.

These challenges have spurred the growth of ‘AI for neuroscience’, a field that uses advances in deep learning to model brain function with increasing accuracy^15,16^. Deep neural networks have shown promise in predicting neural responses to sensory and cognitive inputs^17^. However, many of these models remain modality-specific, limiting their ability to mirror how the brain integrates multisensory information ^4,10–12^. As a result, researchers are turning to multimodal deep learning, which learns from visual, linguistic, and auditory streams to model complex brain functions^18–20^. This trend is supported by neuroscience evidence that cortical responses across regions can be jointly modeled within a common representational space ^21,22^. Still, popular models like CLIP (Contrastive Language-Image Pretraining) ^23^—while successful in aligning image and text features—treat perception as a series of static snapshots, falling short of capturing the temporal continuity central to real-world cognition. Because the brain processes stimuli as events unfolding over time, models that incorporate temporal structure, such as video-text alignment, may offer a more biologically plausible and cognitively meaningful framework.

In this study, we ask a fundamental question in cognitive neuroscience: How can we build models that accurately predict whole-brain neural responses to rich, dynamic, and naturalistic experiences? To answer this, we apply a video-text alignment encoding framework, using VALOR (Vision-Audio-Language Omni-peRception) ^24^—a high-performing, open-source model that aligns visual and linguistic features over time—to predict brain responses during movie watching. By analyzing naturalistic fMRI data from the Human Connectome Project (HCP) ^25^, we conducted four experiments (Fig. 1) to show that our model (1) achieves superior predictive accuracy across both sensory and high-order cognitive areas, (2) generalizes robustly to out-of-distribution stimuli (ShortFunMovies dataset), (3) automatically maps semantic dimensions of cortical organization without manual labeling, and (4) reveals predictive coding gradients linked to individual differences in cognitive ability. Together, these findings demonstrate that multimodal, temporally aligned models offer a powerful and ecologically valid approach to understanding how the brain processes complex, real-world information.

**Fig. 1.**
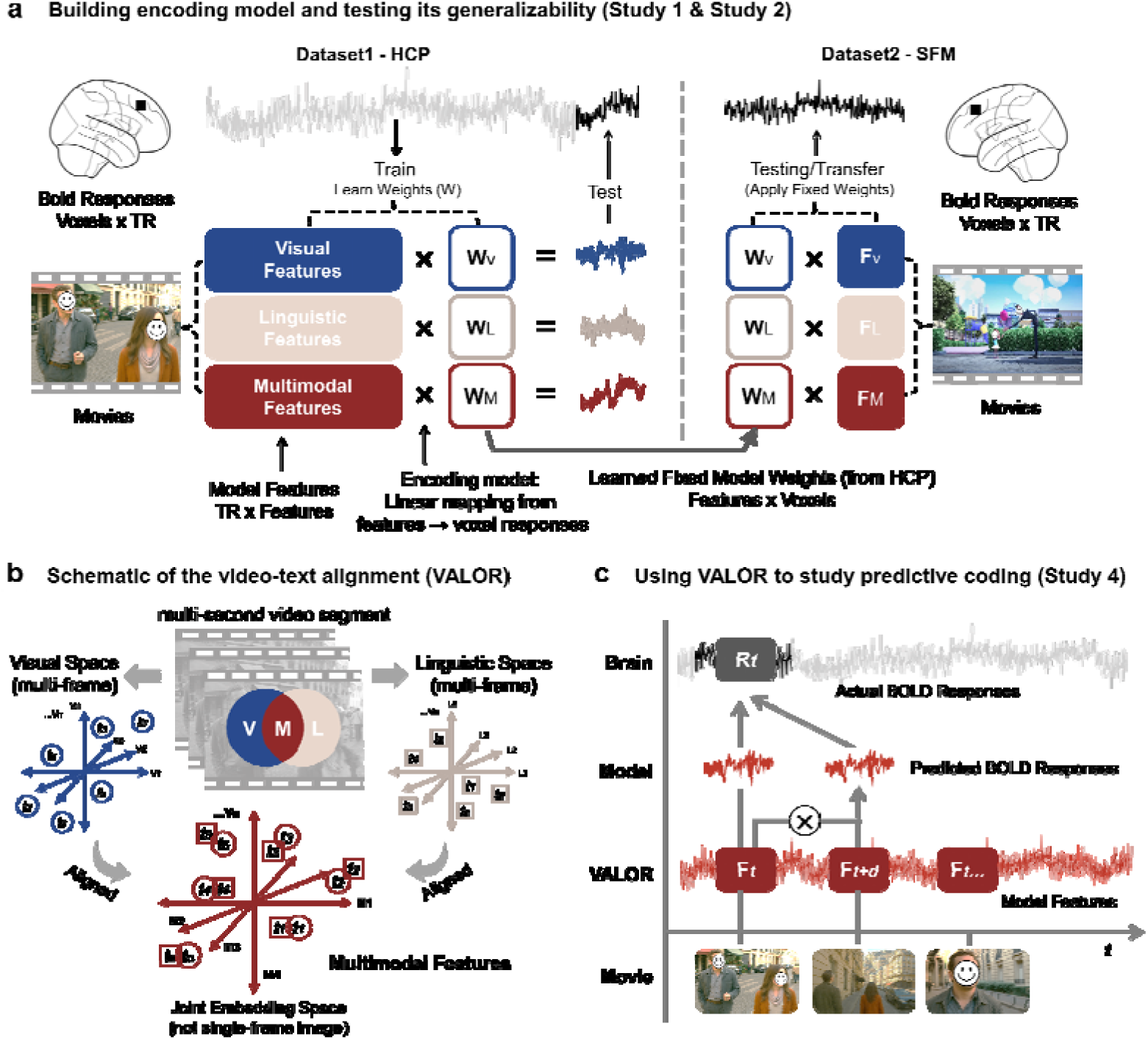
Overview of the Study Design. **a** Encoding model construction and generalization testing (Study 1 & Study 2). Left panel: Whole-brain BOLD responses were recorded while participants viewed naturalistic movies from the HCP 7T dataset. Visual, linguistic, and multimodal (video–text aligned) features were extracted at each TR and used to train voxel-wise encoding models. During training, the encoding model learns voxel-specific weight matrices that map stimulus features to BOLD responses, capturing each voxel’s feature tuning. Right panel: To assess generalization, the learned voxel-wise weights were held fixed and applied to features extracted from novel movies in an independent dataset (ShortFunMovies, SFM), collected at a different site with different participants. Predicted BOLD responses were compared with empirically measured group-mean fMRI responses in the SFM dataset using Pearson correlation, providing a test of cross-subject and cross-dataset transfer of stimulus-locked representations. **b** Schematic of the video–text alignment model (VALOR). VALOR encodes temporally extended video segments (comprising multiple consecutive frames over several seconds) and their associated textual descriptions into separate visual and linguistic embedding spaces. These representations are aligned into a shared 512-dimensional joint embedding space via contrastive learning, such that matched video–text pairs are pulled together while mismatched pairs are pushed apart. By operating on dynamic video input rather than isolated frames, VALOR captures both semantic content and temporal structure, distinguishing it from static image–text alignment models such as CLIP. **c** Using VALOR to study predictive coding (Study 4). To probe predictive coding, the encoding model was extended to include a temporal forecast window. Features at the current time point (F_t_) were combined with features from a future time point (F_t+d_), where d denotes the prediction distance in TRs. Comparing models with and without future features allowed us to quantify regional differences in predictive timescales across cortex. All human-related images in this figure are AI-generated illustrations. No real faces or identifiable individuals are shown.

## Results

### Study 1: Enhanced whole-brain encoding with video-text alignment

In Study 1, we tested whether video-text alignment features offer superior whole-brain neural encoding compared to unimodal and image-based multimodal approaches. We analyzed high-resolution 7T fMRI data from 178 participants in the Human Connectome Project (HCP) as they passively watched one hour of movie stimuli. The movie segments were split into training and test sets, and voxel-wise encoding models were built using kernel ridge regression ^26^ to predict brain responses from four types of input features: (1) AlexNet (visual features), (2) WordNet (semantic features), (3) CLIP (image-based multimodal features), and (4) VALOR, a video-text alignment model that encodes multimodal features from continuous video sequences (Fig. 1b).

As shown in Fig. 2a (left panel), each model exhibited domain-specific strengths: AlexNet performed best in early visual regions, while WordNet, CLIP, and VALOR achieved higher accuracy in language-related and high-level association areas. However, VALOR stood out by achieving significantly higher whole-brain prediction accuracy than all other models (all ps < .04, FDR corrected). To quantify these effects, we computed average prediction accuracy within three sets of regions of interest (ROIs): (1) Visual cortex including V1 and V4, (2) Language-related regions including middle temporal gyrus (MTG) and angular gyrus (AG), and (3) High-level association areas including precuneus (PCu), posterior cingulate cortex (PCC) and medial prefrontal cortex (mPFC).

**Fig. 2.**
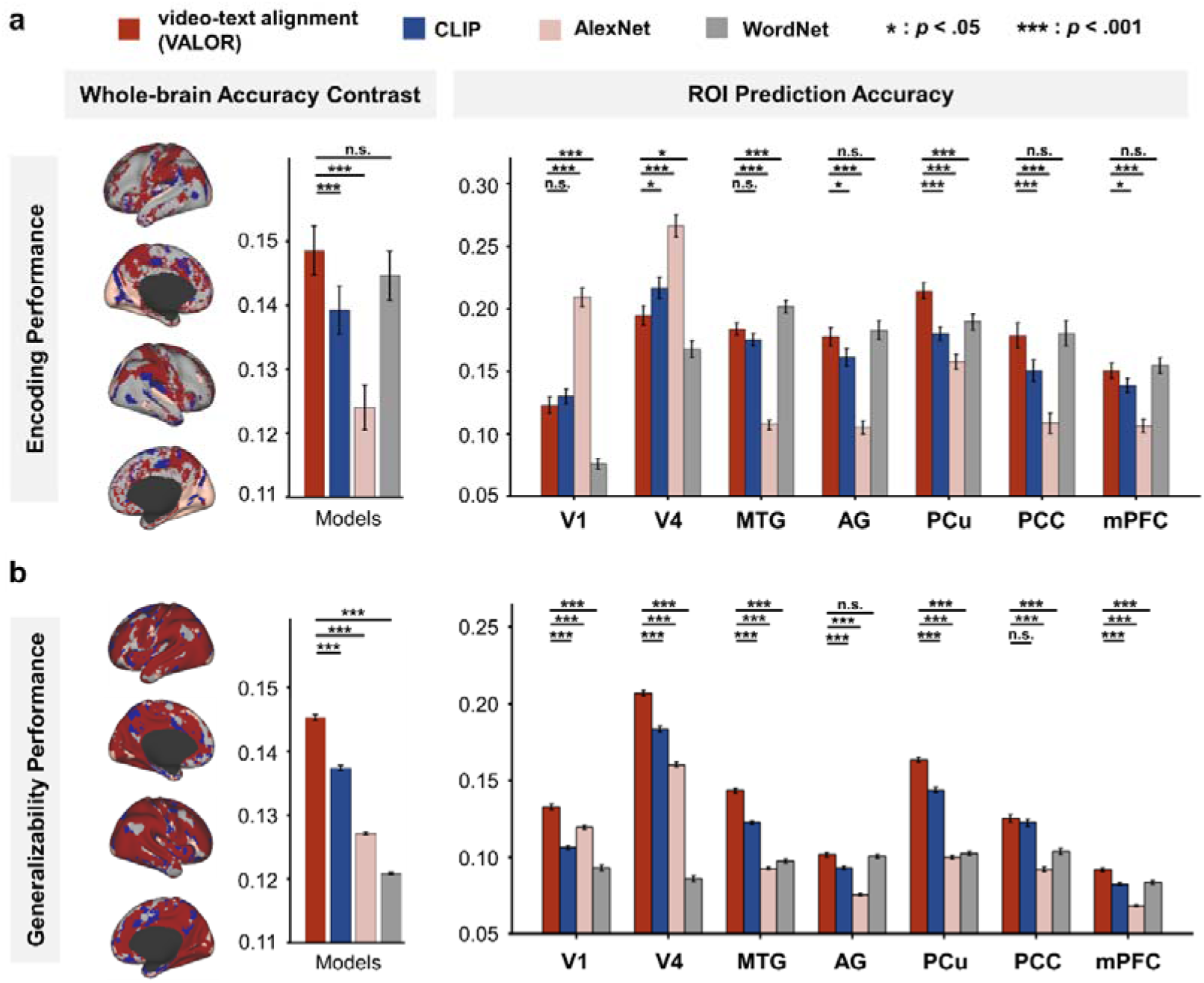
Comparison of encoding and generalization performance across models. **a** Whole-brain encoding performance (Study 1). Voxel-wise and ROI-level comparisons of four encoding models trained on the HCP 7T dataset: VALOR (video-text alignment, red), AlexNet (visual features, blue), WordNet (semantic features, pink), and CLIP (image-based multimodal features, gray). Left panel: Cortical surface maps show the best-performing model at each voxel. Mean whole-brain prediction accuracy across participants, with VALOR significantly outperforming all baselines. Right panel: ROI-wise prediction accuracy across predefined regions (e.g., visual, language, high-level association areas). Bars indicate group-level means, with error bars denoting standard error of the mean (SEM). Statistical comparisons (two-sided paired t-tests, FDR-corrected) are reported only between VALOR and each of the other models. **b** Generalization performance on independent data (Study 2). Models trained on HCP data were tested on the independent ShortFunMovies (SFM) dataset to assess generalizability. Left: Voxel-wise surface maps highlight where VALOR outperforms other models across cortex. Right: ROI-level generalization performance, with color coding and statistical tests consistent with panel (a). For both voxel-wise and ROI analyses, VALOR demonstrates broad generalization across both sensory and high-level regions, exceeding the performance of all other models. For the display of larger brain rendering, please check the Supplementary material (Figure S4)

As shown in Fig. 2a (right panel), AlexNet outperformed others in visual regions, while WordNet and the multimodal models performed better in language and association areas. Importantly, VALOR matched WordNet in language and high-level regions but surpassed CLIP across multiple ROIs, particularly in the precuneus and PCC. Moreover, VALOR showed significantly higher accuracy in visual cortex than WordNet (all ps < .001), demonstrating its balanced predictive coverage across both sensory and cognitive domains. These results were further supported by representational similarity analysis, which revealed stronger correspondence between VALOR features and actual brain activity across ROIs (see Supplementary Fig. S1).

Together, these findings demonstrate that integrating visual and linguistic information over time yields more accurate and comprehensive neural predictions than static or unimodal models. The video-text alignment approach captures the temporal and semantic continuity of naturalistic stimuli, offering a powerful framework for modeling whole-brain activity and advancing our understanding of how the brain processes complex, dynamic experiences.

### Study 2: Robust cross-dataset generalization through video-text alignment

Generalizability is a key benchmark for neural encoding models, reflecting their capacity to predict brain responses to unseen, out-of-distribution stimuli. To evaluate this, we tested whether the video-text alignment model trained on the HCP dataset could generalize to an independent, qualitatively distinct dataset. Specifically, we used our in-house ShortFunMovies (SFM) dataset, in which 20 participants viewed eight short films—six animated and two live-action—differing substantially in style and content from the HCP movies. Full dataset details and analysis procedures are provided in the Methods.

As shown in Fig. 2b (left panel), voxel-wise analysis revealed that VALOR (video-text alignment) significantly outperformed all baseline models—including AlexNet (visual), WordNet (linguistic), and CLIP (image-based multimodal)—across much of the cortex. ROI-level analysis confirmed this pattern: VALOR consistently achieved higher prediction accuracy across nearly all unimodal and multimodal brain regions (Fig. 2b, right panel). These results underscore the importance of temporally integrated multimodal representations for capturing generalizable brain responses.

Although this analysis is not a classical within-subject out-of-distribution generalization test, it evaluates the extent to which different feature spaces capture stimulus-locked representations that are consistent across subjects and transferable across datasets, stimuli, and acquisition environments. Notably, VALOR achieved this without requiring manual preprocessing steps such as principal component analysis (PCA) or semantic annotation, unlike unimodal models. This highlights its strength as a scalable, automated, and ecologically valid framework for modeling brain activity in response to diverse, real-world stimuli.

### Study 3: Comparing data-driven multimodal representations with category-based semantic annotation

A central question in naturalistic encoding is how data-driven feature representations derived from pretrained models relate to more interpretable, category-based semantic annotations that have historically been used to study cortical semantic organization. Although recent work has shown that pretrained language and vision–language models can capture semantic structure without explicit annotation, category-based approaches such as WordNet remain valuable as interpretable reference frameworks. Here, we leverage the WordNet-based semantic components reported by Huth et al. (2012) ^5^ not as a state-of-the-art alternative, but as a historically grounded benchmark, allowing a controlled comparison between data-driven multimodal representations and manually defined semantic categories under matched naturalistic movie stimuli.

To test whether our model could automatically derive similar semantic structures, we applied the video-text alignment encoding model to fMRI data from Huth et al. (2012), in which five participants watched two hours of naturalistic movie clips. First, we assessed voxel-wise prediction accuracy using our multimodal features. As shown in Fig. 3a (and Supplementary Fig. S2), the video-text alignment model consistently outperformed the original WordNet-based model across most of the cortex. Next, we applied principal component analysis (PCA) to encoding model weights, following the approach of Huth et al. (2012). Projecting stimulus features onto each principal component (PC) revealed interpretable semantic dimensions, while projecting encoding weights showed how each PC was represented across the cortex. Analysis of over 20,000 video clips from VALOR’s training set revealed meaningful semantic axes. For instance, PC1 distinguished mobile vs. static content, PC2 separated social vs. non-social categories, PC3 contrasted mechanical vs. non-mechanical stimuli, and PC4 differentiated natural vs. civilizational themes (Fig. 3b). These components closely mirrored those reported in the original WordNet-based model. To quantify this similarity, we computed the Jaccard index between cortical projections of the top four PCs from our model and those derived from WordNet features. As shown in Fig. 3c, we observed substantial spatial overlaps, indicating that our model recovered the semantic organization of human brain comparable to human-labeled ground truth—without any manual annotation.

**Fig. 3.**
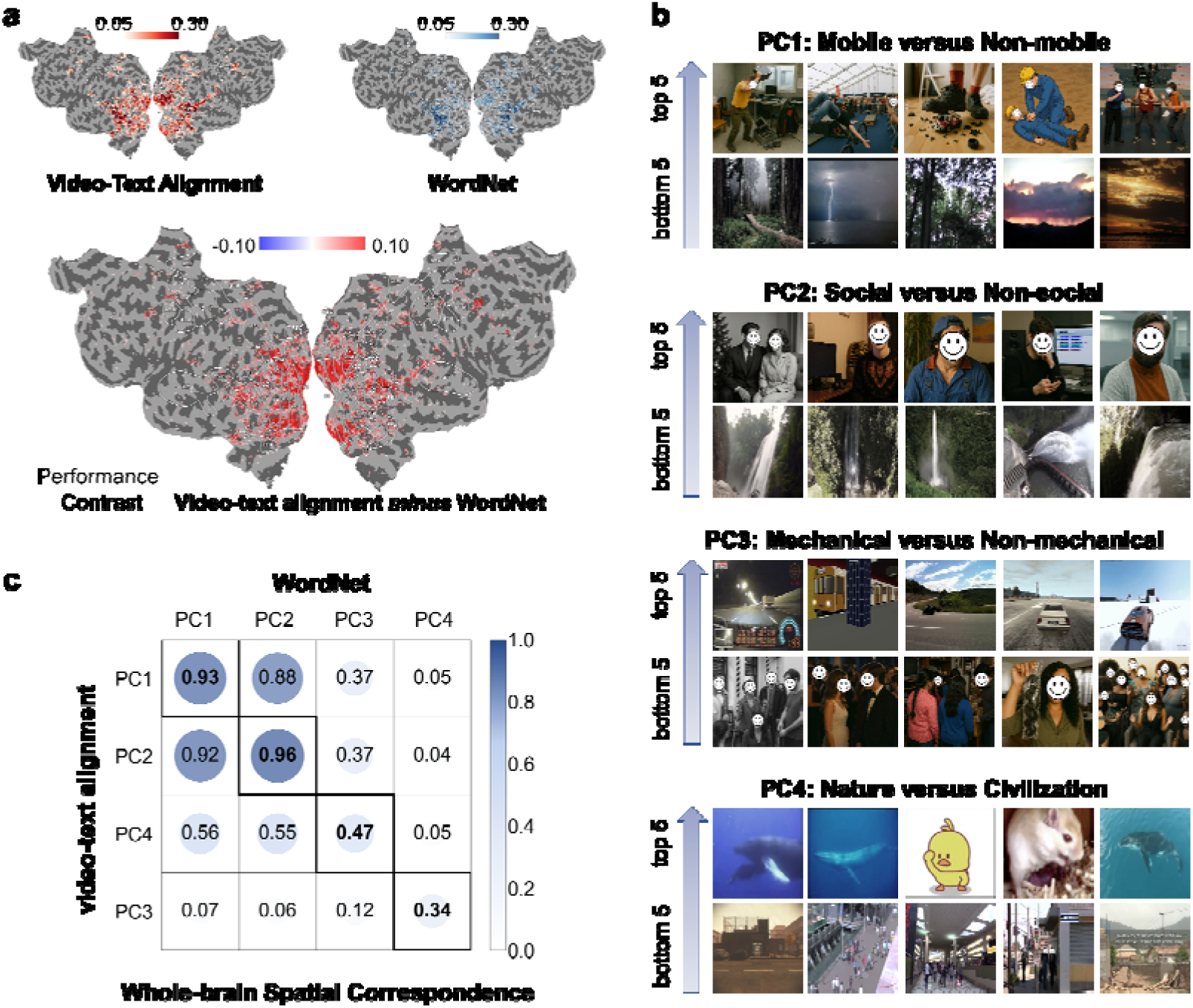
Cortical semantic mapping from video-text alignment features. **a** Voxel-wise prediction performance. Top: Encoding accuracy maps based on video-text alignment features (VALOR) versus WordNet-based semantic features from Huth et al. (2012). Bottom: Difference map showing voxel-wise performance gains (VALOR minus WordNet). Red areas indicate regions where VALOR outperforms WordNet. Results are shown for one representative subject; similar patterns were observed in the remaining four participants (see Supplementary Fig. S2). **b** Semantic dimensions revealed by video-text alignment. Principal component analysis (PCA) was applied to encoding weights derived from VALOR features across ∼20,000 video clips. Example frames illustrate the semantic meaning of the top four PCs: PC1=Mobility (e.g., movement vs. stillness; aligns with Huth’s PC1), PC2=Social content (e.g., people interaction vs. nature; aligns with Huth’s PC2), PC3=Mechanical vs. non-mechanical stimuli (aligns with Huth’s PC4), PC4=Civilization vs. natural environments (aligns with Huth’s PC3). All human-related images in this figure have been replaced with AI-generated illustrations to avoid including identifiable individuals. No real faces or photographs of people are shown. **c** Spatial correspondence with manually annotated semantic maps in Huth et al, (2012). Jaccard similarity matrix quantifying spatial overlap between the top four PCs from the VALOR model (rows) and those from the WordNet-based model (columns). The highest similarity scores for each column were diagonal, indicating strong alignment between automatically derived and manually annotated semantic components. Detailed projections of each semantic component onto the cortical surface can be found in the Supplementary Material (Fig. S3).

Together, these results demonstrate that the video-text alignment encoding model can automatically uncover meaningful, brain-wide semantic structures from naturalistic stimuli. This provides a powerful and scalable alternative to manual labeling, offering new opportunities to study semantic representations in the human brain with greater efficiency and ecological validity.

### Study 4 Neural predictive coding mechanisms through video-text alignment

Predictive processing is a core principle of brain function by which the brain interprets and interacts with the world through actively anticipating future events and continuously updating its internal model to minimize prediction errors^27^. While much of the existing evidence for predictive coding has come from tightly controlled experimental settings^28–30^, its manifestation during naturalistic experiences remains less understood.

To investigate predictive coding in an ecologically valid context, we used the HCP 7T fMRI dataset, focusing on runs where participants viewed Hollywood films—narratives that naturally engage anticipation and inference. We then tested whether incorporating representations of upcoming events—using a “forecast window”—could improve the video-text alignment encoding model’s prediction of neural responses (Fig. 1c). Specifically, for each time point, we combined the current feature representation (F*_t_*) with that of a future segment (F*_t_*_+*d*_), where d indicates the prediction distance ^31^. Comparing models with and without future information yielded “predictive scores”, which quantify the neural benefit of forward-looking representations.

As shown in Fig. 4a, several regions exhibited significant predictive enhancements, including the superior temporal gyrus (STG), middle temporal gyrus (MTG), superior parietal gyrus (SPG), and precuneus (PCu). These findings suggest that the brain actively anticipates upcoming content during movie viewing. To test whether different brain areas operate on distinct predictive timescales, we calculated a “prediction distance” metric for each voxel. Averaging these distances across four ROIs revealed a hierarchical gradient: the STG showed shorter-range predictions, while the MTG, SPG, and especially the PCu anticipated further into the future (Fig. 4b). This pattern aligns with theories of cortical hierarchy^32–34^, suggesting that higher-order regions integrate information over longer temporal windows.

**Fig. 4.**
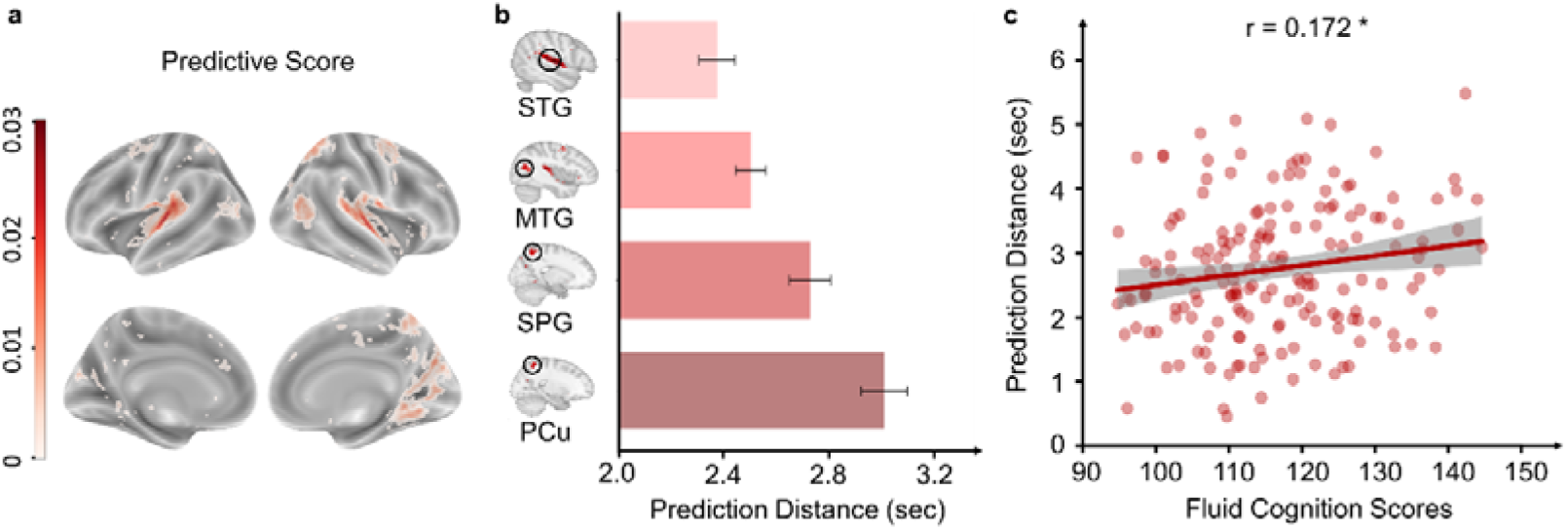
Predictive coding revealed by video-text alignment features. **a** voxel-wise predictive scores. For each voxel, a predictive score was computed as the improvement in prediction accuracy when future stimulus features (F*_t_*_+*d*_) were added to current features (F*_t_*) in the encoding model. Only voxels with significant predictive enhancement across participants are shown (Wilcoxon rank-sum test, FDR-corrected). **b** regional prediction distances. Voxels that showed highest predictive scores were STG, MTG, SPG and PCu. We calculated prediction distance for these regions on a per-voxel, per-participant basis, and then averaged the values. **c** brain–behavior correlation. Scatter plot showing a positive correlation between prediction distance in the SPG and individual fluid cognitive scores across HCP participants (r = 0.172, p < .05, FDR-corrected). This suggests that individuals with broader predictive horizons in the parietal cortex tend to exhibit stronger fluid reasoning ability.

Finally, we examined whether prediction horizons were linked to individual differences in cognition. We focused on fluid intelligence (gF) because gF is widely taken to index domain-general capacities such as maintaining and updating relational context over several seconds, integrating multiple constraints, and exerting flexible top-down control — functions that should support anticipating what will happen next in a continuous narrative. We targeted two parietal regions, the SPG and the PCu, which have both been repeatedly linked to gF and high-level cognitive control in the individual-differences literature^35,36^. For each participant, we correlated fluid cognition scores with that participant’s average prediction horizon in each region. As shown in Fig. 4c, individuals with longer prediction horizons in SPG showed higher fluid cognition scores (SPG: r = 0.172, FDR-corrected p = 0.047). PCu showed a similar positive trend (PCu: r = 0.111, FDR-corrected p = 0.146) but did not reach significance. These associations suggest that the ability to sustain a longer predictive timescale during naturalistic perception co-varies with broader fluid cognitive capacity. No additional brain regions or behavioral measures were examined in this analysis.

In summary, these results show that the video-text alignment model not only captures real-time brain responses to ongoing stimuli but also reveals how the brain projects forward in time, supporting hierarchical prediction and linking anticipatory processing to individual cognitive abilities.

## Discussion

This study introduces video-text alignment encoding as a powerful and ecologically valid framework for modeling whole-brain responses to naturalistic stimuli. By integrating visual and linguistic features over time, our approach addresses long-standing limitations in traditional encoding models and offers key advances across four dimensions: predictive accuracy (Study 1), cross-dataset generalization (Study 2), semantic space mapping (Study 3), and predictive coding mechanisms (Study 4). Collectively, these findings demonstrate that temporally aligned multimodal deep learning can uncover how the brain processes complex, dynamic information in real-world contexts.

In Study 1, we show that video-text alignment models outperform both unimodal and static multimodal approaches in predicting cortical activity (Fig. 2a). While AlexNet and WordNet capture localized visual or linguistic responses, respectively, they fail to generalize beyond their domains. In contrast, VALOR—which fuses visual and linguistic information over time—achieves high prediction accuracy across both low-level sensory and high-level integrative regions, including the precuneus (PCu), insula, and medial prefrontal cortex^37–39^. VALOR matches WordNet in language regions (e.g., angular gyrus) and significantly outperforms it in visual areas (e.g., V1, V4), underscoring that semantic models alone are insufficient to capture naturalistic perception. Additionally, VALOR exceeds the performance of CLIP, a leading static multimodal model, as its training objective aligns multi-second video–text units, enforcing a temporal integration window and event-level semantics that maintain cross-frame consistency and narrative context, whereas CLIP’s image-level alignment provides no intrinsic mechanism for such temporal continuity ^40^. More broadly, this difference reflects distinct inductive biases in how the two models represent visual–linguistic information. CLIP is optimized for framewise image–text correspondence, encouraging representations that emphasize instantaneous visual semantics but remain agnostic to temporal structure. In contrast, VALOR is explicitly biased toward aggregating information over multiple consecutive frames and aligning representations at the level of temporally extended events. These inductive biases favor context maintenance, semantic stabilization, and narrative coherence over time, which are known to be critical for driving responses in association cortex during continuous movie perception. Interestingly, both VALOR and CLIP showed unexpectedly high accuracy in traditionally unimodal sensory areas (e.g., certain voxels in cuneus and lingual gyrus, see Fig. 2a, left panel), suggesting that even early visual cortex may integrate multimodal information during dynamic, real-world experiences ^41,42^. This conclusion is further supported by representational similarity analysis (Supplementary Fig. S1), which shows VALOR’s feature space aligns more closely with measured brain activity than all other models. Together, the relative gains over AlexNet (purely visual), WordNet (manual semantic annotation), and CLIP (static image–text alignment) indicate cortical systems whose responses are best captured by multi-second, multimodal integration, and highlight regions that accumulate and stabilize narrative context over time. At the same time, these findings do not imply that visual–text alignment in the brain gives rise to fully amodal, modality-independent semantic representations. Instead, we suggest that alignment between visual and linguistic signals may serve to stabilize and coordinate scene-level meaning across modalities and over time. From this perspective, VALOR’s architecture—by integrating visual information over multi-second windows and aligning temporally extended video segments with language—provides a computational proxy for how the brain may use linguistic constraints to organize, disambiguate, and maintain coherent representations of unfolding events. The observed encoding gains therefore highlight regions engaged in temporally stabilized, cross-modal integration during naturalistic perception, rather than providing evidence for abstract semantic codes divorced from sensory input.

Beyond predictive accuracy, Study 2 highlights the ecological validity of video-text alignment models by demonstrating their robust generalization to novel stimuli (Fig. 2b). Traditional unimodal models (e.g., AlexNet, WordNet) require manual preprocessing (e.g., PCA, annotation alignment) to adapt to new datasets, while static image-based multimodal models (e.g., CLIP) lack temporal structure, limiting their ability to capture dynamic neural responses. By contrast, VALOR exhibited stronger generalization in a cross-cohort, cross-stimulus, and cross-site transfer evaluation. Together, these findings underscore the importance of distinguishing encoding models that primarily capture shared, stimulus-driven neural structure from those whose performance relies more heavily on subject-specific heterogeneity, particularly when evaluating generalization across participants and datasets. Consistent with this distinction, integrating visual and semantic information over time appears to support more stable and flexible neural representations. This aligns with the idea that the brain continuously adapts to ever-changing environments by integrating multimodal cues across time ^10,43^. By demonstrating robust performance across different datasets, our approach moves beyond static encoding models toward a framework that more closely reflects the brain’s natural adaptability and predictive processing, advancing the ‘AI for neuroscience’ field toward more biologically plausible models of real-world cognition ^44–47^.

Study 3 demonstrates the utility of video–text alignment models for probing higher-order semantic representations during naturalistic perception. Our comparison between VALOR-derived representations and WordNet-based semantic components highlights an important distinction between data-driven and category-based approaches to modeling meaning in the brain. While multimodal pretrained models offer flexible, high-dimensional representations that capture semantic structure without explicit annotation, category-based frameworks provide interpretability and theoretical anchoring ^4,48^. Using WordNet-based labeling from prior work as an interpretable reference point, we show that VALOR automatically extracts semantic dimensions—including mobility, sociality, and civilization—that closely mirror those identified using manual semantic categories (Fig. 3). The observed alignment between VALOR PCs and WordNet semantic components suggests that large-scale semantic organization emerges consistently across these approaches, even though they differ in how semantic structure is defined and learned. This convergence supports the use of pretrained multimodal models as practical encoding tools for naturalistic stimuli, while also underscoring the continued value of interpretable semantic benchmarks for understanding which aspects of meaning are represented across cortex. We do not argue that semantic annotation is required for modern encoding models; rather, WordNet-based features serve here as a historically grounded and interpretable reference for contextualizing data-driven multimodal representations.

In Study 4, we used a video-text alignment model to investigate predictive coding mechanisms. Because our analysis incorporates only future-aligned features, the reported effects should be interpreted as reflecting forward-oriented representations rather than generic sensitivity to temporal context; directly contrasting past- and future-aligned features will be an important direction for future work. By incorporating a forecast window, we found that different brain regions encode future events over distinct timescales (Fig. 4a–b). Short-term predictions were strongest in the superior temporal gyrus (STG), while longer-range forecasts emerged in regions such as the precuneus (PCu), consistent with theories of cortical hierarchy ^32–34,49^. Notably, we observed that individuals with longer prediction distances in the superior parietal gyrus (SPG) exhibited higher fluid cognition scores. This finding suggests a potential link between anticipatory processing and individual cognitive capacity, offering new insights into how the brain predicts and organizes unfolding information in complex environments.

Despite these advances, several limitations remain. First, our analyses relied on a single multimodal architecture (VALOR), operated primarily on TR-level features, and were validated in part on a relatively small independent cohort (n = 20); future work should test whether the same principles hold across alternative models, longer adaptive temporal windows, and larger, more diverse samples. Second, we did not directly compare VALOR to state-of-the-art video-only spatiotemporal models (e.g., Video Swin Transformer, VideoMAE, and related architectures) that are designed to capture temporal visual structure without language grounding; such comparisons will be important for isolating the specific contributions of temporal visual processing versus cross-modal video–text alignment in naturalistic neural responses. Third, for comparability across models we evaluated each model using its single best-performing layer within a matched encoding pipeline rather than using multilayer fusion or ensembling, which allowed us to attribute performance differences to representational format but likely underestimates the absolute performance ceiling. Finally, Finally, part of VALOR’s advantage may reflect model capacity: larger pretrained models often yield higher encoding accuracy, so repeating these analyses with size-matched image-only and image–text models will be critical for disentangling model scale from representational content.

In conclusion, this work establishes video-text alignment encoding as a robust, scalable, and biologically informed framework for studying the brain’s response to naturalistic stimuli. By capturing the temporal and semantic richness of real-world input, this approach advances the field beyond static, modality-limited models and provides new tools for investigating semantic cognition, predictive processing, and individual differences. Beyond theoretical insights, our framework holds practical promise for applications in brain-computer interfaces, clinical neuroimaging, and the development of next-generation cognitive models. As encoding models evolve, video-text alignment stands out as a crucial bridge between deep learning and naturalistic neuroscience, bringing us closer to a comprehensive understanding of the brain in action.

## Methods

### HCP Naturalistic fMRI Dataset

We analyzed high-resolution 7T fMRI data from 178 individuals who participated in the HCP movie-watching protocol. The dataset included four audiovisual movie scans (1 hour in total) with varying content, from Hollywood film clips to independent Vimeo videos. The fMRI data underwent preprocessing using the HCP pipeline, which included correction for motion and distortion, high-pass filtering, regression of head motion effects using the Friston 24-parameter model, removal of artifactual time series identified with ICA, and registration to the MNI template space. Further details on data acquisition and preprocessing can be found in previous publications^25,50^. For our analysis, we excluded rest periods and the first 20 seconds of each movie segment, resulting in approximately 50 minutes of audiovisual stimulation data paired with the corresponding fMRI response.

### Movie Feature Extraction

1. Video–text alignment features (VALOR): To extract video-based multimodal features, we used VALOR (VALOR-large checkpoint), an open-source pretrained video–text alignment model^24^. VALOR combines visual encoders (CLIP and Video Swin Transformer) for extracting visual features and a text encoder (BERT) for extracting textual features ^23,51,52^. These representations are aligned in a shared embedding space through contrastive learning. We segmented each movie at the TR level and, for each segment, extracted VALOR’s projected video–text embedding from the final projection head of the alignment module to obtain a 512-dimensional feature vector. These embeddings were then time-aligned to the corresponding BOLD responses.
2. CLIP features: To compare with static image-based multimodal models, we utilized CLIP (ViT-B/32), which aligns visual and textual representations through contrastive learning but processes individual frames independently without capturing temporal context. One video frame was sampled per TR, and the pooled image embedding after CLIP’s projection into the shared image–text space was extracted to obtain a 512-dimensional feature vector. These TR-aligned vectors were used directly as regressors in the voxel-wise encoding models.
3. AlexNet features: Visual features were extracted by sampling frames at the TR level and processing them with AlexNet, an eight-layer convolutional neural network comprising five convolutional layers followed by three fully connected layers. Features from all five convolutional layers were evaluated in preliminary analyses; the fifth convolutional layer showed the best performance and was used in subsequent analyses. Intra-image z-score normalization was applied to reduce amplitude effects. Principal component analysis (PCA) was used to reduce dimensionality, retaining the top 512 components to match the dimensionality of multimodal feature spaces. This pipeline was implemented using the DNNBrain toolkit ^53^.
4. WordNet features: Semantic features were obtained from publicly available WordNet annotations provided with the HCP dataset (7T_movie_resources/WordNetFeatures.hdf5), following the procedure of Huth et al. (2012). Throughout this manuscript, we use the term “semantic features” to refer to such human-annotated, category-based representations of scene content, and we reserve the term “linguistic features” for continuous language embeddings derived automatically from pretrained language or vision–language models. Each second of the movie clips was manually annotated with WordNet categories according to predefined guidelines: (a) identifying clear objects and actions in the scene; (b) labeling categories that dominated for more than half of the segment duration; and (c) using specific category labels rather than general ones. A semantic feature matrix was constructed with rows corresponding to time points and columns to semantic categories, with category presence coded as binary values. More specific categories from the WordNet hierarchy were added to each labeled category, yielding a total of 859 semantic features. These features were used directly as regressors. We also evaluated a PCA-reduced 512-dimensional variant (fit within each training fold to avoid leakage); because this version performed slightly worse, we report results from the full 859-dimensional representation in the main text. For the generalization analysis in Study 2, annotations for the SFM dataset were aligned to the same WordNet category space to ensure consistency.

These processes described above were performed on a compute node equipped with two Intel(R) Xeon(R) Platinum 8383C CPUs and three NVIDIA GeForce RTX 4090 GPUs. To comply with the journal’s policy, we replaced all human-related video frames used for illustrative purposes (e.g., in Fig. 1 and Fig. 3) with AI-generated images that do not depict real individuals. These synthetic images were created solely for explanatory visualization and were not used in any model training, testing, or analysis.

### Voxel-wise encoding models

Voxel-wise encoding models were created to establish connections between different stimuli and the corresponding brain responses. These models were developed for each individual using visual, linguistic, and multimodal features, and kernel ridge regression was used to map them to brain activation while preserving the interpretability of the model weights. The training set consisted of all movie segments from the HCP 7T movie dataset, excluding repeated segments, while the test set consisted of the repeated segments to improve noise reduction and reliability by averaging the brain imaging data over four runs per participant. A finite impulse response model with four delays (2, 4, 6, and 8 seconds) was used to account for the hemodynamic response. The regularization parameters for each voxel and subject were optimized through 10-fold cross-validation, exploring 30 parameters ranging logarithmically from 10 to 10^30^. Model performance was assessed on the test data by calculating Pearson correlation coefficients between predicted and observed voxel activation sequences.

### Novel movie dataset for generalizability test

To evaluate the generalizability of encoding models trained on the HCP dataset, we collected a new fMRI dataset—referred to as “ShortFunMovies”(SFM)—from 20 adult female Chinese participants (mean age = 23.76 ± 2.26 years). This dataset offers exceptional stimulus diversity through eight different movie segments (45 mins in total), including six animations and two live actions, that differ markedly in content and style from the HCP movie stimuli. Each participant watched these eight audiovisual movies while undergoing an fMRI scan using a 3 Tesla Siemens Prisma Magnetom scanner at the MRI Center of Beijing Normal University. The scanning parameters were as follows: TR = 2000 ms, TE = 30 ms, flip angle = 90°, FOV = 210 × 210 mm, spatial resolution = 2.5 mm^³^, and multiband factor = 6. After preprocessing the fMRI data with fMRIPrep, we denoised and smoothed the brain imaging data (fwhm = 6 mm). We then extracted visual, linguistic, image-based and video-based multimodal features from the movie stimuli and fed them into subject-specific encoding models that were pretrained on the HCP dataset. These models were used to predict each individual’s brain activation patterns in response to the new stimuli. To assess the generalizability of the models, we calculated Pearson correlation coefficients between the predicted brain activations of each HCP subject models and the actual group-level mean activation observed in participants exposed to the new movie dataset. This new data collection was approved by the Institutional Review Board of Beijing Normal University (IRB_A_0024_2021002), and informed consent was obtained from all participants. All participants received monetary compensation after completing the MRI scanning. The SFM dataset was deposited in the OSF website.

### Semantic space analysis

Whole-brain semantic space analysis has traditionally relied on manual stimulus annotation. Huth et al. (2012) created a semantic feature matrix by manually annotating video content and decoding semantic dimensions across the brain. To address this challenge, we wanted to verify whether our video-text alignment model could accurately and efficiently automate the analysis of whole-brain semantic space. Therefore, we repeated Huth et al.’s analysis and compared our results with theirs.

We first fit voxel-wise encoding models using VALOR features for each of the five participants in the Huth et al. dataset. For each participant, this yielded a weight map linking each VALOR feature to each voxel. We then stacked these weight maps across participants to form a single voxel-by-feature weight matrix and applied principal component analysis (PCA).

The top four principal components from this analysis (“VALOR PCs”) captured shared spatial patterns of VALOR feature weights across cortex. To interpret these components, we projected VALOR feature vectors from >20,000 video segments in the VALOR training set onto each PC, which revealed dominant semantic axes (e.g., mobility, sociality, civilization). For comparison, we used the semantic principal components reported by Huth et al. (2012) from their WordNet-based encoding model; these “WordNet PCs” were taken directly from the published work and were not refit or reweighted using VALOR. To analyze how PCs from WordNet and video-text alignment features are distributed in the cortex, we projected their weights onto the cortical surfaces of subjects. By calculating Jaccard coefficients for the PC distributions in both encoding models, we measured variations and similarities across different cortical regions. The overall Jaccard index, obtained by averaging these coefficients across all participants, provides a comprehensive view of the shared neural patterns.

### Probing Predictive Coding Mechanisms

To test predictive coding when watching movies, we incorporated forecast representations— so-called ‘forecast window’ — and examined whether this significantly improved the prediction accuracy of video-text alignment encoding models across different voxels^31^. These forecast windows, denoted as *F_t_*_+*d*_, include features extracted by the video-text alignment model from n-second clips, where n aligns with the TR duration of one second. The end frame of each segment is aligned with a temporal offset *d* from the current TR. *F_t_*_+*d*_ contains information from a segment subsequent to the current TR, reflecting the brain’s prediction mechanism for potential future stimuli encounters.

For each subject *s* and voxel *v*, we calculated the ‘Brain score’ *B*^(*s*,*v*)^, reflecting the effectiveness of using features from the current TR clip to predict brain activation:

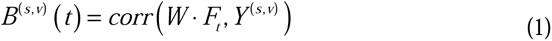

with *W* as the kernel ridge regression, *F_t_* as the feature of the current TR clip, *corr* as Pearson’s correlation and *Y*^(*s*,*v*)^ as the fMRI signals of one individual *s* at one voxel *v* .

For each prediction distance *d*, subject *s*, and voxel *v*, we computed the ‘Brain score’ *B*_(*d*,*s*,*v*)_, reflecting the model’s accuracy when it includes features of the forecast window alongside the present TR clip features:

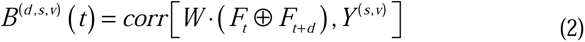

where *F_t_*⊕*F_t_*_+*d*_ represents the integrated feature set combining the current and forecast clip features.

The ‘Predictive score’ *P*_(*d*,*s*,*v*)_ was computed as the improvement in brain score when concatenating forecast windows to present multimodal features:

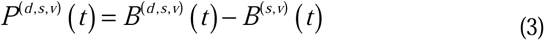

To ensure dimensionality compatibility between *F_t_* ⊕ *F_t_*_+*d*_, we applied PCA for dimensionality reduction, reducing both feature types to 100 dimensions each.

We defined the optimal ‘prediction distance’ for each individual *s* and voxel *v* as:

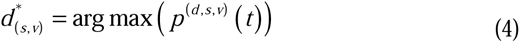

This process utilized the HCP movie-watching dataset, specifically focusing on the second and fourth runs due to their Hollywood film content, which typically evokes continual plot conjectures during viewing, and regarded the last movie segment of run 4 as the test set and other segments as the training set.

## Data and Code availability

All data and analysis codes in this project were uploaded to Open Science Framework (https://osf.io/2fnr4/)

## Acknowledgement

This work was supported by National Natural Science Foundation of China (32422033, 32171032, 32430041), National Science and Technology Innovation 2030 Major Program (2022ZD0211000, 2021ZD0200500), Open Research Fund of the State Key Laboratory of Cognitive Neuroscience and Learning (CNLZD2103) and the start-up funding from the State Key Laboratory of Cognitive Neuroscience and Learning, IDG/McGovern Institute for Brain Research, Beijing Normal University (to Y.W.)

## Author contributions

M.F. and Y.W. conceived and designed the research. M.F., G.C., Y.Z., and M.Z. collected and analyzed the data. M.F., G.C., Y.Z., and M.Z. wrote the initial draft of the manuscript. M.F. and Y.W. edited and reviewed the final manuscript.

## Supplementary Materials

### Representational Similarity Analysis (RSA) in Study 1

To assess the relationship between video-text alignment features and neural activity prior to constructing the encoding model, we conducted a representational similarity analysis (RSA). This approach allowed us to evaluate how well different feature types—AlexNet (visual), WordNet (linguistic), CLIP (image-based multimodal), and VALOR (video-based multimodal)—aligned with multivoxel brain activation patterns across key cortical regions. We focused on three groups of functionally defined regions of interest (ROIs): (i) Visual regions including V1 and V4, (ii) Language-associated regions including middle temporal gyrus (MTG) and angular gyrus (AG), (iii) Higher-order cognitive regions including medial prefrontal cortex (mPFC), posterior cingulate cortex (PCC), and precuneus (PCu)

Each HCP movie run was split into 4–5 stimulus segments that exhibited consistency in visual or linguistic content. Each segment was analyzed independently using a multivariate RSA framework. We computed representational dissimilarity matrices (RDMs) based on pairwise Pearson correlations between multivoxel patterns across timepoints (TRs), yielding TR-by-TR matrices for each ROI. Corresponding feature-based RDMs were constructed for each model using the extracted features. To account for the hemodynamic delay, the first four TRs of each run were excluded. We then computed feature-brain RSA similarity by correlating each brain RDM with the feature RDM using Spearman’s rank correlation. Analyses were performed at the individual subject level and results were aggregated across participants for group-level comparisons. We used paired-sample t-tests with FDR correction to assess statistical differences in RSA values between models.

Figure S1 summarizes the results. In early visual cortex (V1), VALOR’s RSA performance matched that of AlexNet and exceeded that of WordNet and CLIP. In V4, VALOR outperformed all other models. In language areas (MTG and AG), VALOR showed substantially higher RSA values than both unimodal and static multimodal features. This advantage also extended to high-level cognitive regions (PCu, PCC), where VALOR consistently yielded the highest RSA scores. The mPFC was the only region where VALOR and WordNet features performed comparably.

These results underscore the broad representational fidelity of video-text alignment features across visual, language, and integrative networks. By capturing both semantic and temporal structure, VALOR provides a more comprehensive approximation of real neural representations, supporting its use in modeling whole-brain responses to naturalistic stimuli.

**Fig. S1.**
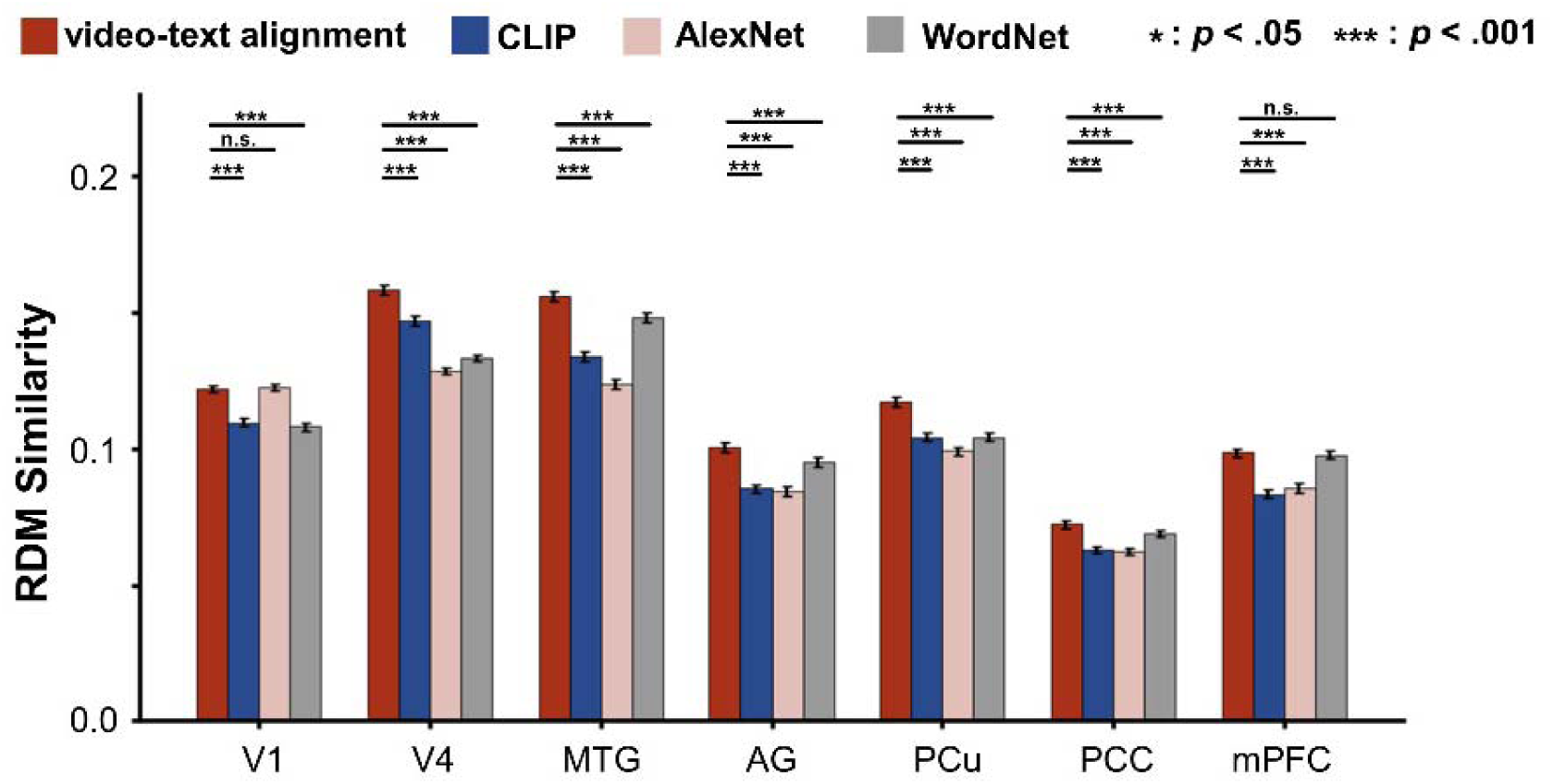
Representational similarity analysis (RSA) across feature types and brain regions. Bar plots show RSA results comparing four feature extraction models—video-text alignment (VALOR), CLIP, AlexNet, and WordNet—across seven predefined regions of interest (ROIs): V1, V4 (visual), MTG, AG (language), and PCu, PCC, mPFC (higher-order cognitive regions). RSA values reflect the similarity between multivoxel neural patterns and feature-derived representational dissimilarity matrices (RDMs), aggregated across all participants and movie segments from the HCP 7T dataset. VALOR consistently outperformed or matched other models across most ROIs, particularly in V4, MTG, AG, PCu, and PCC, indicating its ability to capture both low-level perceptual and high-level semantic structure. In V1, VALOR was comparable to AlexNet. In mPFC, performance was similar between VALOR and WordNet. Statistical comparisons were conducted using paired-sample t-tests, with false discovery rate (FDR) correction applied for multiple comparisons. Asterisks indicate significance levels (*p < .05, ***p < .001); "n.s." = not significant. Error bars represent ±1 standard error of the mean (SEM) across participants.

**Figure S2.**
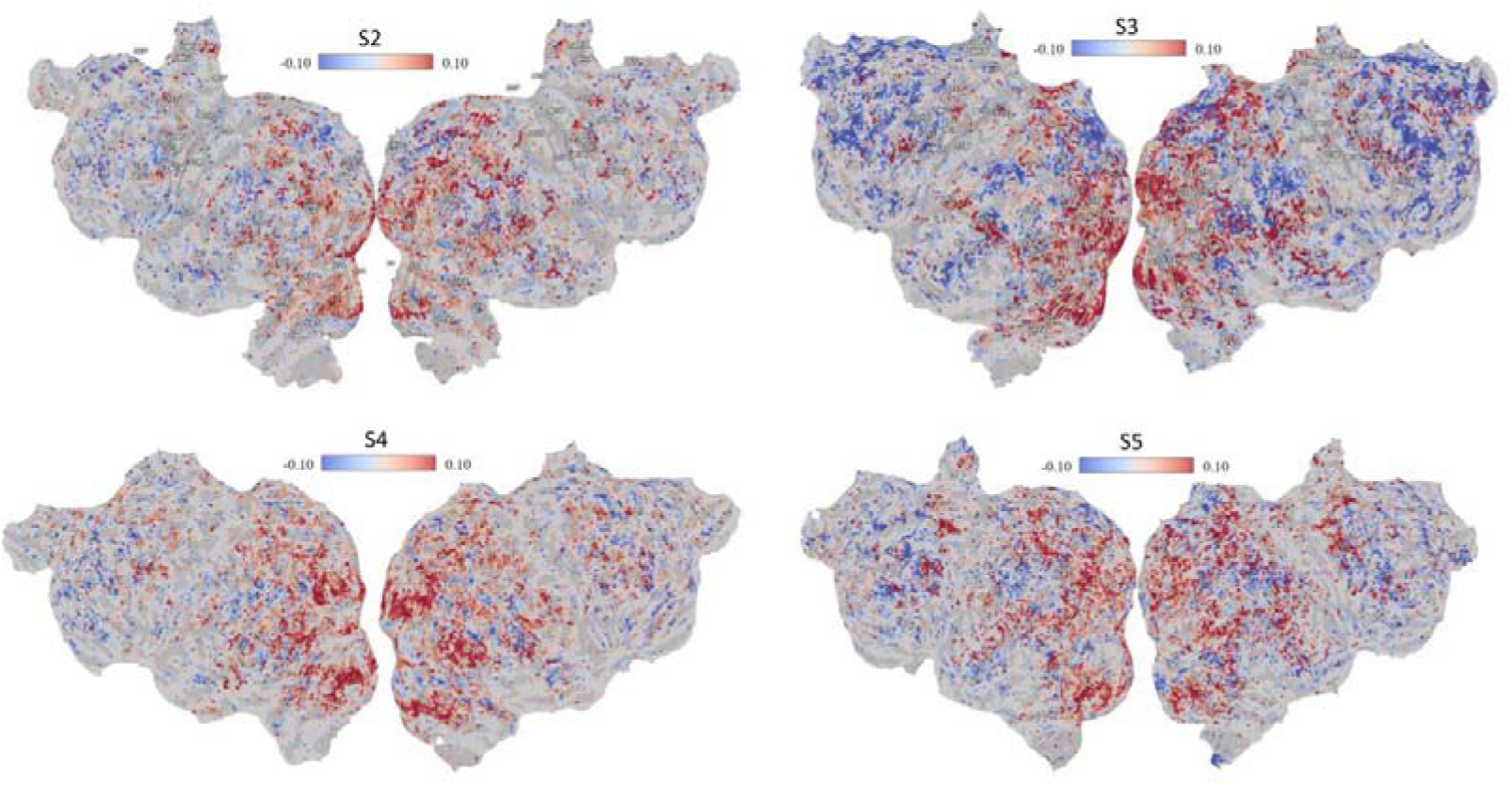
Individual subject maps showing performance difference between feature models (related to Figure 3a). Cortical flat maps show voxel-wise differences in prediction accuracy between the video-text alignment model (VALOR) and the WordNet-based semantic model, computed as VALOR minus WordNet. Data are shown for four individual subjects (S2–S5) from the Huth et al. (2012) dataset. Red voxels indicate regions where VALOR achieved higher prediction accuracy than WordNet; blue voxels reflect regions where WordNet outperformed VALOR. These maps mirror the analysis presented in Figure 3a (main text) and demonstrate that VALOR consistently outperforms WordNet across a wide range of cortical areas, including both hemispheres and across subjects. The color scale reflects the magnitude of performance differences, ranging from -0.10 to 0.10 in Pearson correlation units.

**Figure S3.**
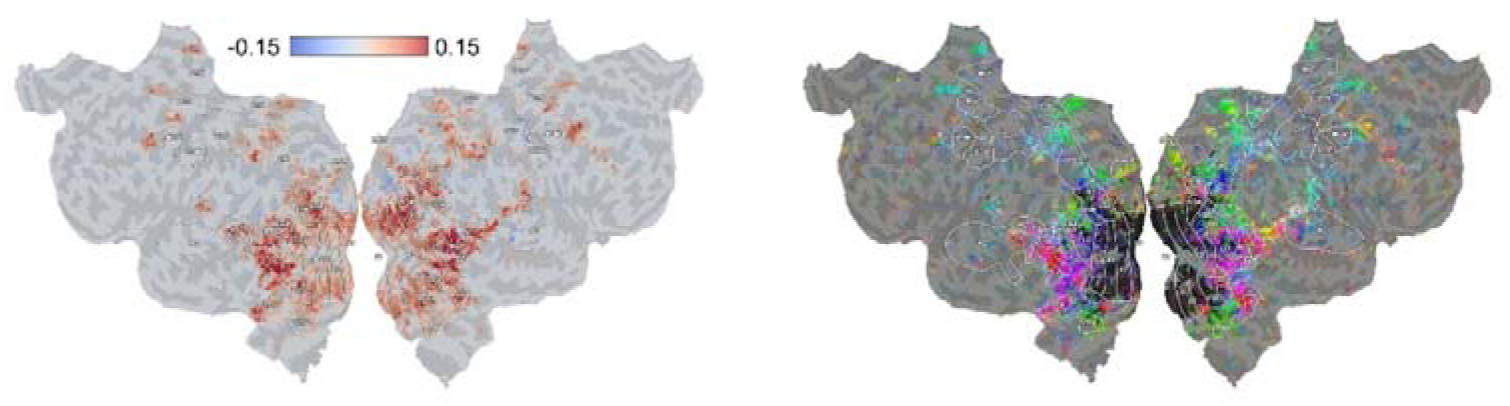
Projections of semantic components onto the cortical surface. Left: Component 1 (PC1) loadings projected onto the individual surface of S1 as continuous, signed weights (color bar indicates magnitude). Right: Components 2–4 displayed with a discrete scheme: PC2 = red, PC3 = green, PC4 = blue (greater saturation denotes larger loadings; near-zero shown in gray).

**Figure S4.**
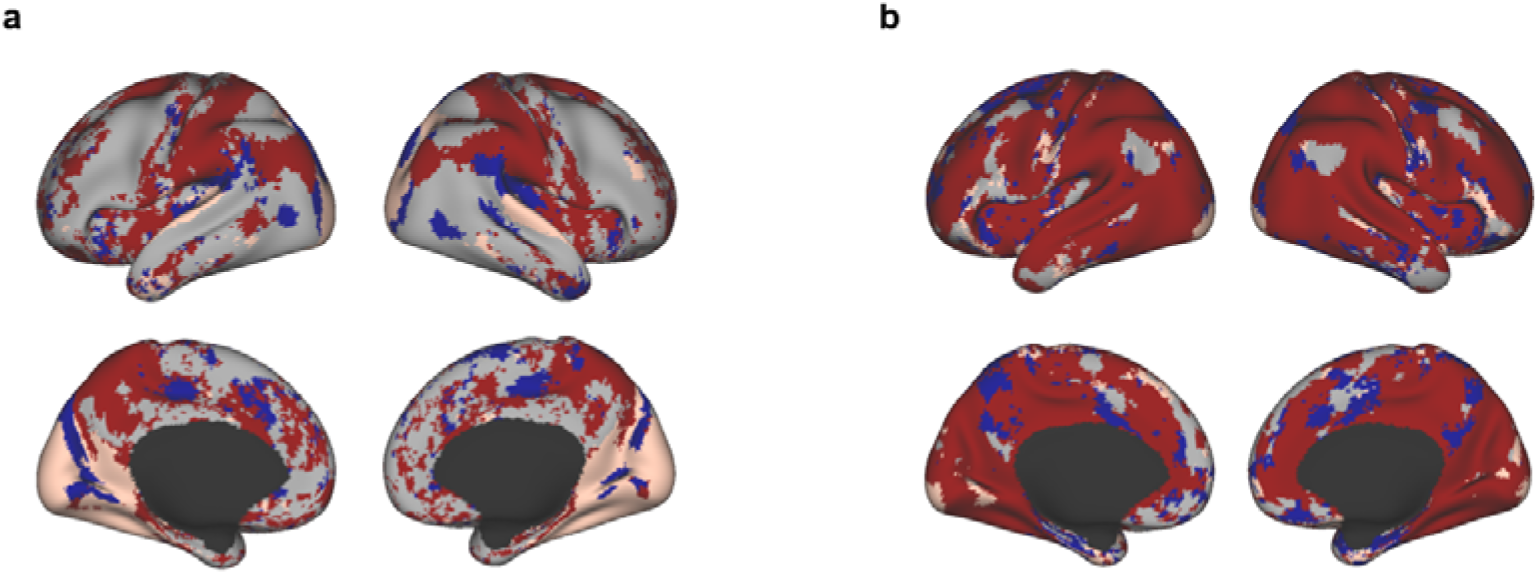
Cortical surface projections of voxel-wise winner-take-all maps showing the best-performing representation at each voxel. (a) Encoding performance on HCP (Study 1). (b) Generalization performance on SFM (Study 2; trained on HCP, tested on SFM). Colors indicate the representation with the highest prediction accuracy: VALOR (red), AlexNet (blue), WordNet (pink), and CLIP (gray).

## References

1. Naselaris, T., Kay, K. N., Nishimoto, S. & Gallant, J. L. Encoding and decoding in fMRI. NeuroImage 56, 400–410 (2011).

2. Tang, J., Du, M., Vo, V. A., Lal, V. & Huth, A. G. Brain encoding models based on multimodal transformers can transfer across language and vision. Preprint at http://arxiv.org/abs/2305.12248 (2023).

3. Wang, A. Y., Kay, K., Naselaris, T., Tarr, M. J. & Wehbe, L. Better models of human high-level visual cortex emerge from natural language supervision with a large and diverse dataset. Nat. Mach. Intell. https://doi.org/10.1038/s42256-023-00753-y (2023) doi:10.1038/s42256-023-00753-y.

4. Huth, A. G., Nishimoto, S., Vu, A. T. & Gallant, J. L. A Continuous Semantic Space Describes the Representation of Thousands of Object and Action Categories across the Human Brain. Neuron 76, 1210–1224 (2012).

5. Huth, A. G., De Heer, W. A., Griffiths, T. L., Theunissen, F. E. & Gallant, J. L. Natural speech reveals the semantic maps that tile human cerebral cortex. Nature 532, 453–458 (2016).

6. Finn, E. S. Is it time to put rest to rest? Trends Cogn. Sci. 25, 1021–1032 (2021).

7. Simony, E. & Chang, C. Analysis of stimulus-induced brain dynamics during naturalistic paradigms. NeuroImage 216, 116461 (2020).

8. Wen, H., Shi, J., Chen, W. & Liu, Z. Transferring and generalizing deep-learning-based neural encoding models across subjects. NeuroImage 176, 152–163 (2018).

9. Tang, J., LeBel, A., Jain, S. & Huth, A. G. Semantic reconstruction of continuous language from non-invasive brain recordings. Nat. Neurosci. 26, 858–866 (2023).

10. Khosla, M., Ngo, G. H., Jamison, K., Kuceyeski, A. & Sabuncu, M. R. Cortical response to naturalistic stimuli is largely predictable with deep neural networks. Sci. Adv. 7, eabe7547 (2021).

11. Ratan Murty, N. A., Bashivan, P., Abate, A., DiCarlo, J. J. & Kanwisher, N. Computational models of category-selective brain regions enable high-throughput tests of selectivity. Nat. Commun. 12, 5540 (2021).

12. Van Uden, C. E. et al. Modeling Semantic Encoding in a Common Neural Representational Space. Front. Neurosci. 12, 437 (2018).

13. Miller, G. A., Beckwith, R., Fellbaum, C., Gross, D. & Miller, K. J. Introduction to WordNet: An On-line Lexical Database*. Int. J. Lexicogr. 3, 235–244 (1990).

14. Madan, S. et al. Benchmarking Out-of-Distribution Generalization Capabilities of DNN-based Encoding Models for the Ventral Visual Cortex. Preprint at http://arxiv.org/abs/2406.16935 (2024).

15. Akkus, C. et al. Multimodal Deep Learning. Preprint at http://arxiv.org/abs/2301.04856 (2023).

16. Perez-Martin, J. et al. A comprehensive review of the video-to-text problem. Artif. Intell. Rev. 55, 4165–4239 (2022).

17. Yamins, D. L. K. & DiCarlo, J. J. Using goal-driven deep learning models to understand sensory cortex. Nat. Neurosci. 19, 356–365 (2016).

18. Dirani, J. & Pylkkänen, L. The time course of cross-modal representations of conceptual categories. NeuroImage 277, 120254 (2023).

19. Kehl, M. S. et al. Single-neuron representations of odours in the human brain. Nature 634, 626–634 (2024).

20. Van Der Linden, M., Van Turennout, M. & Fernández, G. Category Training Induces Cross-modal Object Representations in the Adult Human Brain. J. Cogn. Neurosci. 23, 1315–1331 (2011).

21. Du, C., Fu, K., Li, J. & He, H. Decoding Visual Neural Representations by Multimodal Learning of Brain-Visual-Linguistic Features. IEEE Trans. Pattern Anal. Mach. Intell. 45, 10760–10777 (2023).

22. Oota, S. R., Arora, J., Rowtula, V., Gupta, M. & Bapi, R. S. Visio-Linguistic Brain Encoding. Preprint at http://arxiv.org/abs/2204.08261 (2022).

23. Radford, A. et al. Learning Transferable Visual Models From Natural Language Supervision.

24. Chen, S., et al. VALOR: Vision-Audio-Language Omni-Perception Pretraining Model and Dataset. Preprint at http://arxiv.org/abs/2304.08345 (2023).

25. Glasser, M. F. et al. The minimal preprocessing pipelines for the Human Connectome Project. NeuroImage 80, 105–124 (2013).

26. Dupré La Tour, T., Eickenberg, M., Nunez-Elizalde, A. O. & Gallant, J. L. Feature-space selection with banded ridge regression. NeuroImage 264, 119728 (2022).

27. Millidge, B., Seth, A. & Buckley, C. L. Predictive Coding: a Theoretical and Experimental Review. Preprint at http://arxiv.org/abs/2107.12979 (2022).

28. Meyer, T. & Olson, C. R. Statistical learning of visual transitions in monkey inferotemporal cortex. Proc. Natl. Acad. Sci. 108, 19401–19406 (2011).

29. Shain, C., Blank, I. A., Van Schijndel, M., Schuler, W. & Fedorenko, E. fMRI reveals language-specific predictive coding during naturalistic sentence comprehension. Neuropsychologia 138, 107307 (2020).

30. Todorovic, A., Van Ede, F., Maris, E. & De Lange, F. P. Prior Expectation Mediates Neural Adaptation to Repeated Sounds in the Auditory Cortex: An MEG Study. J. Neurosci. 31, 9118–9123 (2011).

31. Caucheteux, C., Gramfort, A. & King, J.-R. Evidence of a predictive coding hierarchy in the human brain listening to speech. *Nat*. Hum. Behav. 7, 430–441 (2023).

32. Baldassano, C. et al. Discovering Event Structure in Continuous Narrative Perception and Memory. Neuron 95, 709–721.e5 (2017).

33. Hasson, U., Yang, E., Vallines, I., Heeger, D. J. & Rubin, N. A Hierarchy of Temporal Receptive Windows in Human Cortex. J. Neurosci. 28, 2539–2550 (2008).

34. Raut, R. V., Snyder, A. Z. & Raichle, M. E. Hierarchical dynamics as a macroscopic organizing principle of the human brain. Proc. Natl. Acad. Sci. 117, 20890–20897 (2020).

35. Jung, R. E. & Haier, R. J. The Parieto-Frontal Integration Theory (P-FIT) of intelligence: Converging neuroimaging evidence. Behav. Brain Sci. 30, 135–154 (2007).

36. Preusse, F., Elke, V. D. M., Deshpande, G., Krueger, F. & Wartenburger, I. Fluid Intelligence Allows Flexible Recruitment of the Parieto-Frontal Network in Analogical Reasoning. Front. Hum. Neurosci. 5, (2011).

37. Grall, C., Equita, J. & Finn, E. S. Neural unscrambling of temporal information during a nonlinear narrative. Cereb. Cortex 33, 7001–7014 (2023).

38. Kauttonen, J., Hlushchuk, Y., Jääskeläinen, I. P. & Tikka, P. Brain mechanisms underlying cue-based memorizing during free viewing of movie Memento. NeuroImage 172, 313–325 (2018).

39. Richter, F. R., Cooper, R. A., Bays, P. M. & Simons, J. S. Distinct neural mechanisms underlie the success, precision, and vividness of episodic memory. eLife 5, e18260 (2016).

40. Rasheed, H., Khattak, M. U., Maaz, M., Khan, S. & Khan, F. S. Fine-tuned CLIP Models are Efficient Video Learners. in 2023 IEEE/CVF Conference on Computer Vision and Pattern Recognition (CVPR) 6545–6554 (IEEE, Vancouver, BC, Canada, 2023). doi:10.1109/CVPR52729.2023.00633.

41. Bonnici, H. M., Richter, F. R., Yazar, Y. & Simons, J. S. Multimodal Feature Integration in the Angular Gyrus during Episodic and Semantic Retrieval. J. Neurosci. 36, 5462–5471 (2016).

42. Sepulcre, J., Sabuncu, M. R., Yeo, T. B., Liu, H. & Johnson, K. A. Stepwise Connectivity of the Modal Cortex Reveals the Multimodal Organization of the Human Brain. J. Neurosci. 32, 10649–10661 (2012).

43. Driver, J. & Noesselt, T. Multisensory Interplay Reveals Crossmodal Influences on ‘Sensory-Specific’ Brain Regions, Neural Responses, and Judgments. Neuron 57, 11–23 (2008).

44. Brodersen, K. H. et al. Generative Embedding for Model-Based Classification of fMRI Data. PLoS Comput. Biol. 7, e1002079 (2011).

45. Kriegeskorte, N. & Douglas, P. K. Interpreting encoding and decoding models. Curr. Opin. Neurobiol. 55, 167–179 (2019).

46. Sohn, W., Di, X., Liang, Z., Zhang, Z. & Biswal, B. B. Explorations of using a convolutional neural network to understand brain activations during movie watching. Psychoradiology 4, kkae021 (2024).

47. Suter, G., Černis, E. & Zhang, L. Interpersonal computational modelling of social synchrony in schizophrenia and beyond. Psychoradiology 5, kkaf011 (2025).

48. Finn, E. S. & Bandettini, P. A. Movie-watching outperforms rest for functional connectivity-based prediction of behavior. NeuroImage 235, 117963 (2021).

49. Hasson, U., Chen, J. & Honey, C. J. Hierarchical process memory: memory as an integral component of information processing. Trends Cogn. Sci. 19, 304–313 (2015).

50. T. Vu, A., et al. Tradeoffs in pushing the spatial resolution of fMRI for the 7T Human Connectome Project. NeuroImage 154, 23–32 (2017).

51. Devlin, J., Chang, M.-W., Lee, K. & Toutanova, K. BERT: Pre-training of Deep Bidirectional Transformers for Language Understanding. Preprint at http://arxiv.org/abs/1810.04805 (2019).

52. Liu, Z. et al. Video Swin Transformer. in 2022 IEEE/CVF Conference on Computer Vision and Pattern Recognition (CVPR) 3192–3201 (IEEE, New Orleans, LA, USA, 2022). doi:10.1109/CVPR52688.2022.00320.

53. Chen, X. et al. DNNBrain: A Unifying Toolbox for Mapping Deep Neural Networks and Brains. Front. Comput. Neurosci. 14, 580632 (2020).

